# Dopamine receptor-expressing neurons are differently distributed throughout layers of the motor cortex to control dexterity

**DOI:** 10.1101/2023.08.31.555724

**Authors:** Przemyslaw E. Cieslak, Sylwia Drabik, Anna Gugula, Aleksandra Trenk, Martyna Gorkowska, Kinga Przybylska, Lukasz Szumiec, Grzegorz Kreiner, Jan Rodriguez Parkitna, Anna Blasiak

**Affiliations:** Department of Neurophysiology and Chronobiology, Institute of Zoology and Biomedical Research, Jagiellonian University, Krakow, Poland; Department of Molecular Neuropharmacology, Maj Institute of Pharmacology, Polish Academy of Sciences, Krakow, Poland; Department of Brain Biochemistry, Maj Institute of Pharmacology, Polish Academy of Sciences, Krakow, Poland

## Abstract

The motor cortex comprises the primary descending circuits for flexible control of voluntary movements and is critically involved in motor skill learning. Motor skill learning is impaired in patients with Parkinson’s disease, but the precise mechanisms of motor control and skill learning are still not well understood. Here we have used transgenic mice, electrophysiology, *in situ* hybridization and neural tract-tracing methods to target genetically defined cell types expressing D1 and D2 dopamine receptors (D1+ and D2+, respectively) in the motor cortex. We observed that D1+ and D2+ neurons are organized in highly segregated, non-overlapping populations. Moreover, based on *ex vivo* patch-clamp recordings, we showed that D1+ and D2+ cells have distinct morphological and electrophysiological properties. Finally, we observed that chemogenetic inhibition of D2+, but not D1+ neurons, disrupts skilled forelimb reaching in adult mice. Overall, these results demonstrate that dopamine receptor-expressing cells in the motor cortex are highly segregated and play a specialized role in manual dexterity.

## Introduction

The brain’s dopamine system is critical for movement control and motor learning, as evidenced by its role in Parkinson’s disease (PD), a degenerative, neurological disorder characterized by motor deficits caused by progressive dopamine depletion (1, 2). Primary motor symptoms of PD are tremor, rigidity and bradykinesia, but they are often accompanied by upper limb motor impairments, particularly in fine motor skills, including reaching and grasping (3–6). Movement abnormalities in PD arise from disordered neural activity in the cortico-basal ganglia circuits resulting from the loss of dopamine signaling (7–9). Dopamine action is mediated by G_s_-coupled D1 and G_i_- coupled D2 dopamine receptors and neurons expressing these receptors are highly segregated in the basal ganglia circuits (10–12), which suggests that they have specialized roles in the regulation of motor function. In line with this hypothesis, animal studies utilizing optogenetics and chemogenetics show, that cell type-specific modifications of activity of D1 and D2 receptor-expressing neurons in the basal ganglia modulate specific parkinsonian motor behaviors (13, 14).

Despite extensive research on the function of D1 and D2 receptor-expressing projection neurons in the basal ganglia, still little is known about the role of dopamine receptor-expressing neurons in cortical circuits. In this study, we focused on the primary motor cortex, a command center governing voluntary motor control and performance of fine motor skills (15). In rodents, this cortical region is strongly innervated by dopamine fibers and is enriched in both D1 and D2 receptors (16–18). Importantly, dopamine signaling in the motor cortex is necessary for skill learning and synaptic plasticity, as these processes are disrupted in animal models of PD (19–22). Nevertheless, due to the complexity of motor cortex circuits precise mechanisms of fine motor control are still not well understood. Therefore, using transgenic mice, electrophysiology, *in situ* hybridization, neural tract-tracing and chemogenetics we aimed to determine laminar organization of D1 and D2 receptor-expressing neurons in the motor cortex, their properties and role in skilled forelimb reaching in adult mice.

## Results

### Primary motor cortex D1+ and D2+ neurons are organized in differently distributed, highly separated populations

To determine the distribution of D1 and D2 receptor-expressing neurons (D1+ and D2+, respectively) throughout the layers of the primary motor cortex (M1) we used dopamine receptor-specific reporter lines: Drd1aCre mice injected with DIO-mCherry virus and double transgenic Drd2Cre::Ai14 mice (**Figure 1A**). We counted fluorescently labeled cells throughout the superficial (I, II/III) and deep output (V-VI) layers. We found that D1+ cells were mainly distributed throughout the deep output layers V-VI, while D2+ positive neurons were primarily distributed in superficial layer II/III (**Figure 1B and C**). To further verify if M1 neurons co-express D1 and D2 receptors, we used double transgenic Drd1a-tdTomato::Drd2Cre mice injected with DIO-EYFP virus (**Figure 1D**). We counted fluorescently labeled D1+ (tdTomato-positive) and D2+ (EYFP-positive) cells and found that the majority of studied neurons were positive for only one fluorescent reporter, with only 4,15% positive for both tdTomato and EYFP (**Figure 1E**). Overall, these results indicate that D1+ and D2+ neurons in M1 are organized in differently distributed, highly separated populations.

**Figure 1.**
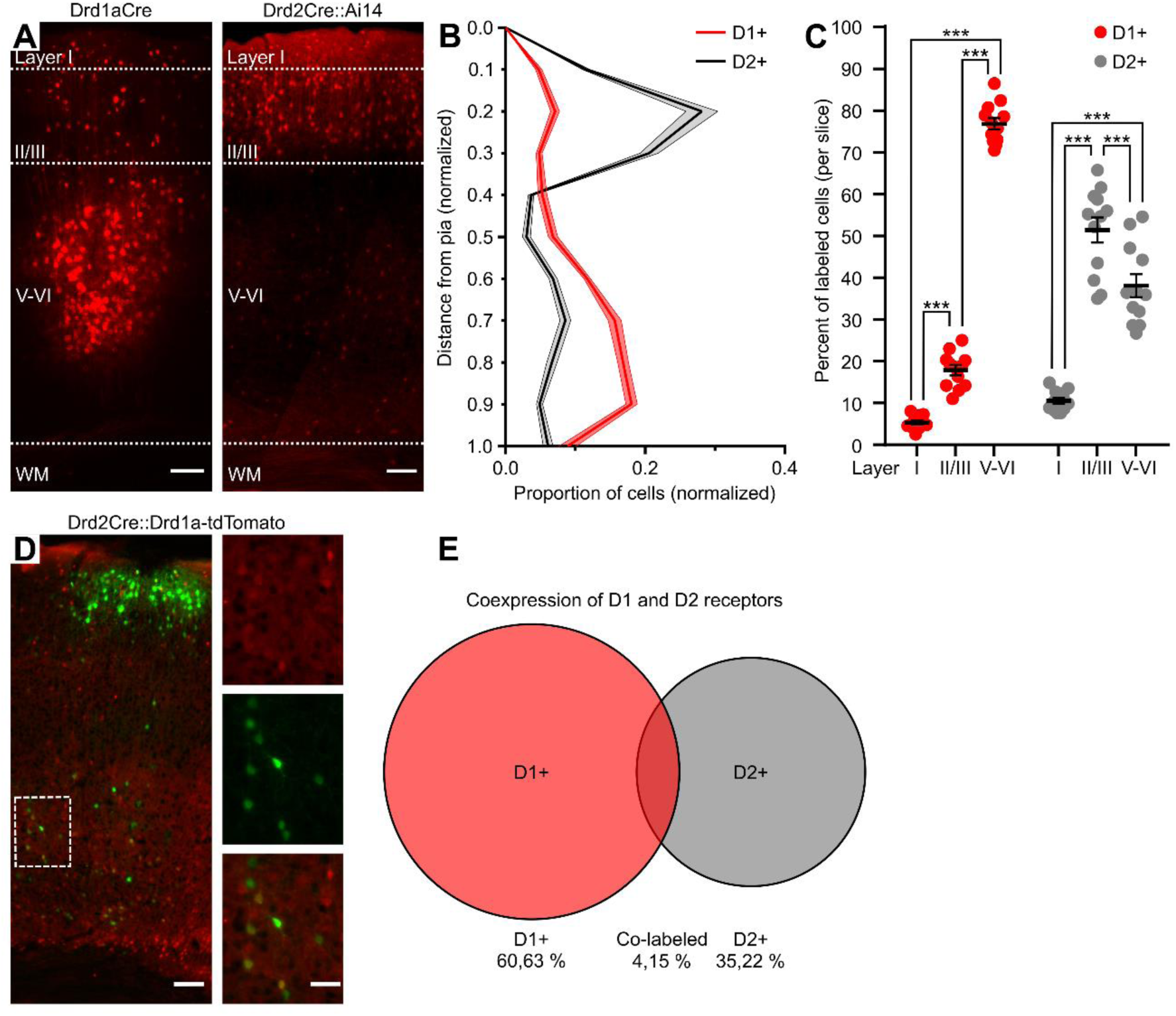
D1+ and D2+ cells are distinctly distributed through layers of the motor cortex. (**A**) Coronal sections obtained from a Drd1aCre mouse injected with DIO-mCherry virus and a Drd2Cre::Ai14 mouse showing laminar distribution of D1+ and D2+ neurons across layers of M1. Laminar boundaries are designated with white dashed lines. WM - white matter. Scale bar: 100 µm (**B** and **C**) Laminar distribution of fluorescently labeled D1+ and D2+ neurons. (**B**) Somatic distance is measured in normalized units (0, pia; 1, white matter), cell number per unit is normalized to a total number of fluorescently labeled cells within the section. (**C**) Distribution in layers I, II/III, V-VI represented as a percentage of fluorescently labeled neurons in all layers. (**D**) A coronal section obtained from a double transgenic Drd1a-tdTomato::Drd2Cre mouse injected with DIO-EYFP virus. Insets show single- (tdTomato or EYFP) and double-labeled cells. Scale bars: main image 100 µm, inset 50 µm. (**E**) Venn diagram showing the overlap between Drd1a (tdTomato-positive) and Drd2 (EYFP-positive) expressing neurons. (**B**, **C** and **E**) Slices were obtained from Drd1aCre, 3 Drd2Cre::Ai14 and 3 Drd1a-tdTomato::Drd2Cre mice (*n* = 3 mice per group, *n* = 4 slices per animal). (**B** and **C**) Results are displayed as mean ± SEM. (**C**) *** *P* < 0.001, 2-way ANOVA followed by Bonferroni’s post hoc test. (**B** and **E**) No statistical tests were applied.

### Distinct neurochemical profiles define the layer-specific distribution of D1+ and D2+ M1 neurons

Cortical neurons can be broadly classified into glutamatergic pyramidal cells or GABAergic interneurons (23). To determine whether D1+ and D2+ neurons in M1 belong to one or the other, RNAscope multiplex *in situ* hybridization was performed, using brain sections from Drd1a-tdTomato and Drd2Cre::Ai14 mice (**Figure 2A**) and specific probes for tdTomato, vesicular glutamate (vGlut2) and GABA (vGAT1) transporters mRNA, as well as D1 and D2 receptors mRNA (**Figure 2B and Supplemental Figure 1**). In both mouse lines, the observed tdTomato mRNA expression pattern corresponded to the distribution of tdTomato-immunolabeled D1+ and D2+ neurons described in the previous section (**Figure 1A**). Furthermore, the majority of tdTomato mRNA expressing (tdTomato+) cells in slices from Drd1a-tdTomato mice were located within deep layers V-VI, while tdTomato+ neurons in slices from Drd2Cre::Ai14 mice were primarily present in layer II/III (**Figure 2A and Supplemental Table 1**). Most of tdTomato+ cells in both Drd1a-tdTomato and Drd2Cre::Ai14 mice co-expressed vGlut2 mRNA (vGlut2+), indicating that they are primarily glutamatergic pyramidal neurons, with the remaining smaller proportion being vGAT1 mRNA expressing (vGAT1+) GABAergic neurons, and a marginal fraction of cells co-expressing mRNA for both transporters (vGlut2+/vGAT1+) (**Figure 2C**). Interestingly, the observed neurochemical phenotypes of tdTomato+ neurons in Drd1a-tdTomato and Drd2Cre::Ai14 animals depended on their laminar distribution (**Figure 2D**). In deep cortical layers V-VI (**Figure 2D right**) most tdTomato+ neurons in Drd1a-tdTomato mice were vGlut2+ and constituted a vast majority of all vGlut2+ neurons identified in this area, while the remaining tdTomato+ cells belonged to a smaller vGAT1+ population. In contrast, the fraction of tdTomato+ neurons co-expressing vGlut2 and vGAT1 mRNA in Drd2Cre::Ai14 mice in the deep layers was almost equal. This pattern was reversed in the superficial layer II/III (**Figure 2D left**), where populations of tdTomato+ cells co-expressing vGlut2 and vGAT1 mRNA in Drd1a-tdTomato mice were more evenly distributed, while the majority of tdTomato+ cells in Drd2Cre::Ai14 mice co-expressed vGlut2 mRNA. Consistent with the results shown in **Figure 1D and E**, we observed minimal *Drd2* and *Drd1a* mRNA co- localization in tdTomato+ cells in Drd1a-tdTomato and Drd2Cre::Ai14 mice, respectively (**Supplemental Figure 1 and Supplemental Table 2**). These results provide further evidence supporting the segregation of D1+ and D2+ neurons in M1 into distinct subpopulations, with D1+ cells located mainly in the deep layers and displaying predominantly glutamatergic phenotype and D2+ neurons being GABAergic, while the opposite is true in the superficial layers.

**Figure 2.**
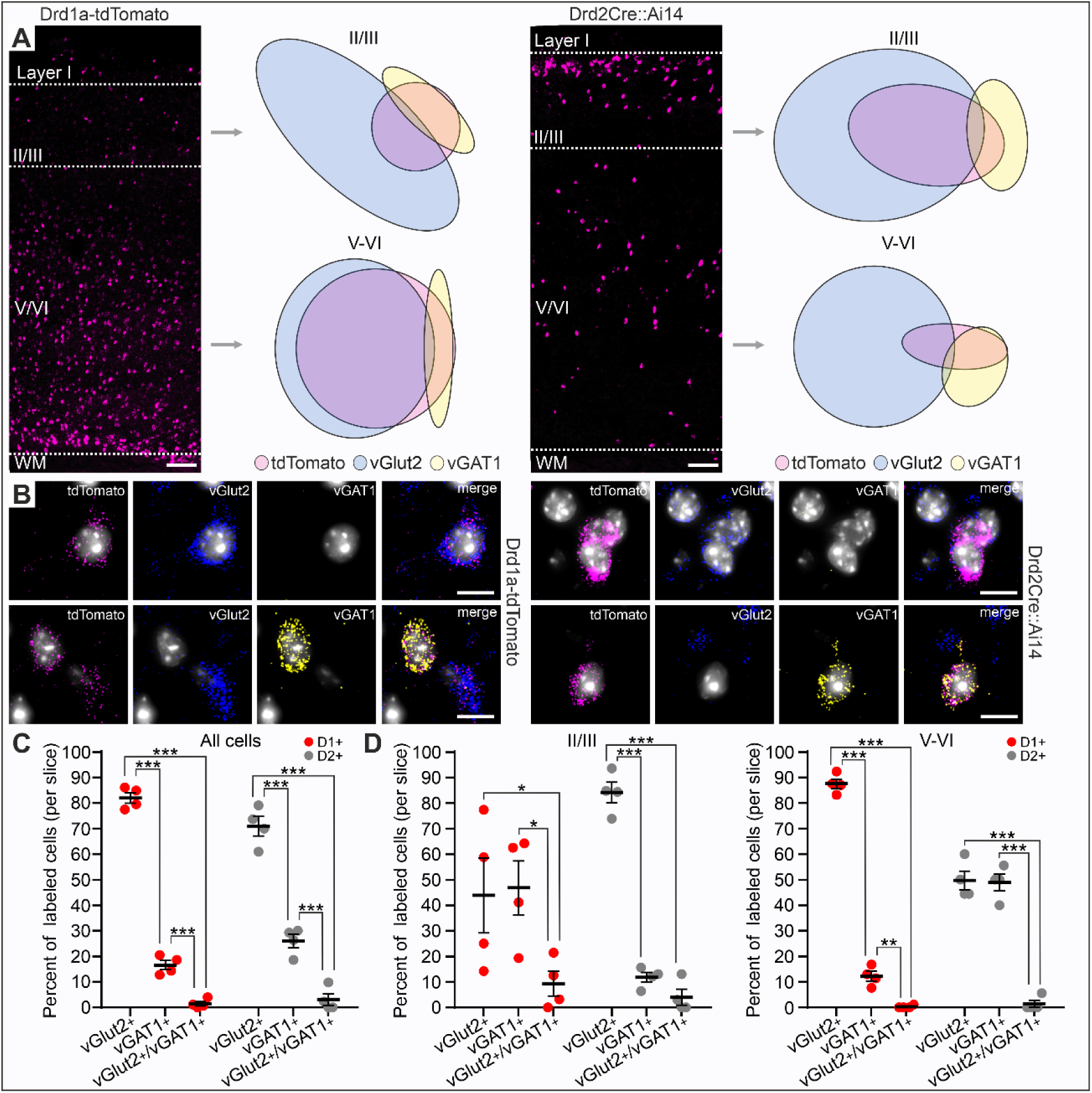
D1+ and D2+ neurons are primarily glutamatergic and GABAergic neuronal populations. (**A**) Coronal sections obtained from a Drd1a-tdTomato and Drd2Cre::Ai14 mouse showing tdTomato mRNA expression across layers of M1. Laminar boundaries are designated with white dashed lines. WM - white matter. Scale bar: 100 µm. Area-proportional Euler diagrams represent proportions and relationships between clusters of neurons identified based on mRNA expression. (**B**) Representative images showing co-localization of tdTomato (pink) mRNA and either vGlut2 (blue) or vGAT1 (yellow) mRNAs within an individual neuron. Scale bar: 10 µm. (**C**) Proportion of neurons co-expressing vGlut2, vGAT1 or mRNAs for both transporters represented as a percentage of all tdTomato+ labeled neurons. (**D**) Proportion of neurons co-expressing vGlut2, vGAT1 or mRNAs for both transporters represented as a percentage of tdTomato+ neurons found in layer II/III (left) or layers V-VI (right). (**C**-**D**) Slices are obtained from Drd1a-tdTomato and Drd2Cre::Ai14 mice (*n* = 2 mice per group, *n* = 2 slices per animal). Results are displayed as mean ± SEM. *** *P* < 0.001, ** *P* < 0.01, * *P* < 0.05, 2-way ANOVA followed by Bonferroni’s post hoc test, with the exception of Euler diagrams, where no statistical tests were applied.

### D1+ and D2+ pyramidal neurons have distinct electrophysiological and morphological properties

Previous studies have shown that D1+ and D2+ layer V pyramidal cell subclasses in the prefrontal cortex have unique electrophysiological and morphological properties (24–26). To test if similar diversity applies to the motor cortex, whole-cell patch clamp recordings and subsequent morphological analysis were performed from fluorescently labeled D1+ and D2+ layer V pyramidal neurons in brain slices obtained from Drd1a-tdTomato and Drd2Cre::Ai14 mice, respectively (**Figure 3A**). To test for possible differences in the voltage dependence of the current flowing through the membrane of D1+ and D2+ neurons, their steady state current-voltage (I-V) relationship was characterized and compared using the linear regression curve slope (**Figure 3C**). The performed analysis showed that examined slopes differed significantly and subsequent post-hoc tests revealed differences between tested groups at low voltage step values, indicating differences in the types or properties of ion channels expressed by D1+ and D2+ neurons. We did not observe differences between D1+ and D2+ neurons in their resting membrane potential or passive membrane properties (resistance, capacitance, time constant) (**Figure 3D**). To characterize and compare the excitability of D1+ and D2+ neurons, the relationship between the injected current and recorded firing rate was evaluated using the linear regression curve slope (**Figure 3E**). The performed analysis revealed that D2+ neurons excitability curve slope was significantly steeper than D1+ neurons, indicating higher excitability (stronger neuronal gain) of D2+ cells.

**Figure 3.**
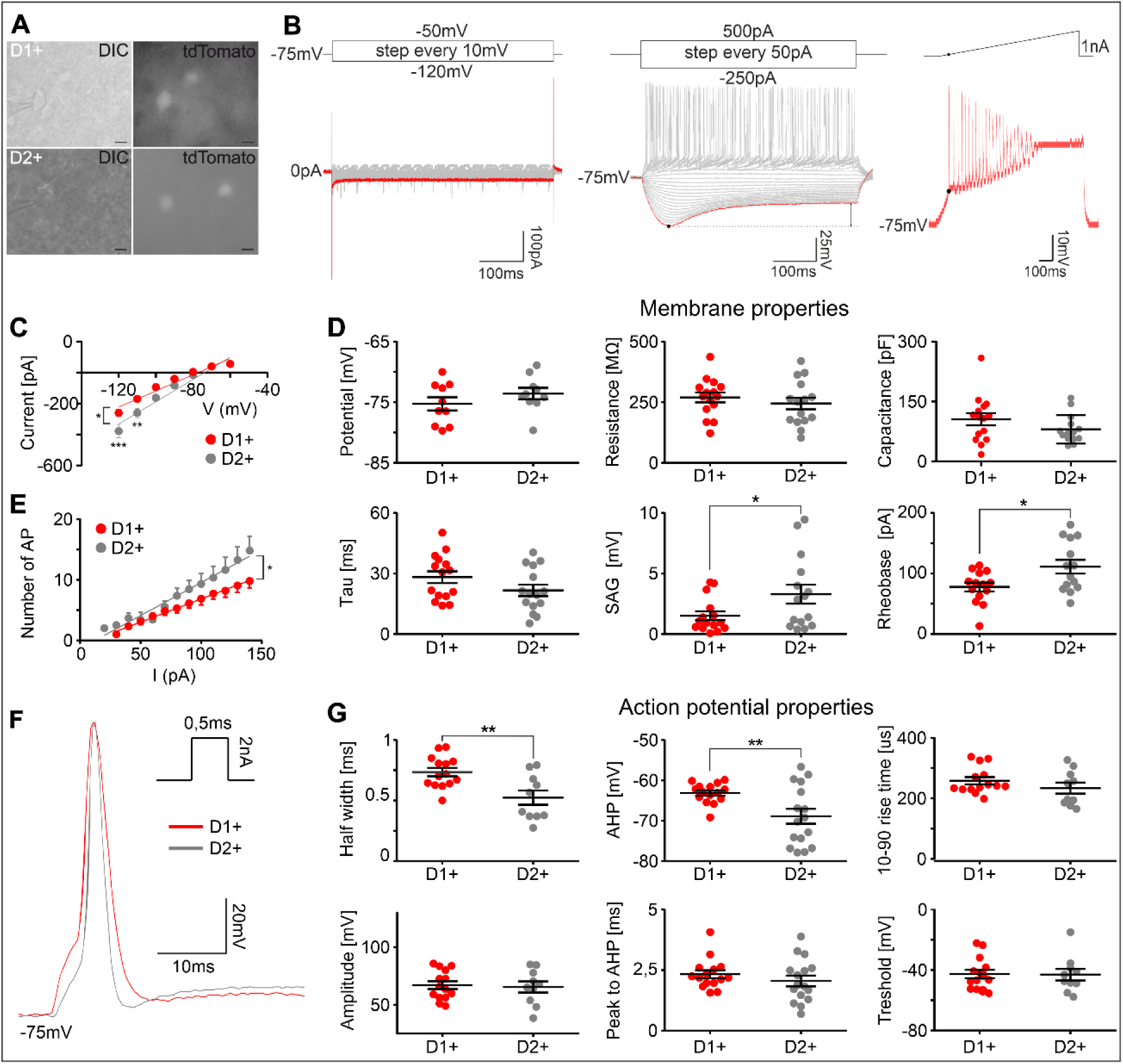
Layer V D1+ and D2+ pyramidal neurons exhibit unique electrophysiological properties. (A) An example of layer V D1+ and D2+ pyramidal neurons recorded by a patch clamp pipette in differential interference contrast (DIC) and fluorescence modes. Slices were obtained from Drd1a-tdTomato (upper panels) and Drd2Cre::Ai14 (lower panels) mice. Scale bars: 10 µm. **(B)** Stimulation protocols (upper panels) and corresponding current responses (bottom panels) of exemplary layer V pyramidal neuron. Voltage (left) and current (middle) steps protocols, and ramp current protocol (right). **(C)** The current- voltage relationship of D1+ and D2+ neurons, measured from the steady-state current responses to voltage step pulses shown in **B** (left panel). D1+ and D2+ neurons displayed highly linear steady state voltage I–V relationship (R^2^ = 0.95 for both groups). (**D**) Membrane properties of D1+ and D2+ neurons. Membrane potential was recorded in 0 current clamp mode. Resistance, capacitance and time constant (tau) were measured from the voltage response to a -50 pA hyperpolarizing current pulse. Voltage sag was measured from the voltage response to a -250 pA hyperpolarizing current step marked in red in **B** (middle panel). (**E**) The input-output (I-O) relationship reflecting the excitability of D1+ and D2+ neurons, quantified by measuring the number of spikes elicited by incremental current pulses (**B**, middle panel). D1+ and D2+ neurons displayed a highly linear I-O relationship (R^2^ = 0.99 for D1+ and 0.97 for D2+). (**F**) D1+ and D2+ neurons exemplary waveforms of a single, evoked AP elicited from membrane potential of -75 mV, with a 0.5 ms rectangle current injection. (**G**) Action potential properties of D1+ and D2+ neurons. (**C-G**) Overall, *n* = 18 D1+ and *n* = 19 D2+ neurons were tested during patch clamp recordings. Results are displayed as mean ± SEM. *** *P* < 0.001, ** *P* < 0.01, * *P* < 0.05, unpaired Student’s t-test or 2-way ANOVA followed by Bonferroni’s post hoc test.

Previous studies showed that D1+ and D2+ prefrontal cortex (PFC) pyramidal neurons can be identified based on the characteristics of the voltage sag induced by hyperpolarization-activated cation currents (I_h_-currents). Specifically, D1+ PFC neurons were characterized by the absence of voltage sag in response to hyperpolarizing current (25), while D2+ neurons were characterized by a large sag amplitude (24, 26). In line with these findings, we observed significantly larger voltage sag amplitude in D2+ than in D1+ M1 pyramidal neurons (**Figure 3D**). Moreover, analysis of the shape of evoked action potentials (AP) revealed, that D2+ M1 pyramidal neurons have smaller AP half width and lower value of afterhyperpolarization (AHP) trough when compared to D1+ cells (**Figure 3F and G**). Importantly, the observed higher sag amplitude and shorter action potential duration in D2+ neurons may contribute to greater excitability of D2+ relative to D1+ neurons (27). At the same time, ramp current stimulations revealed that rheobase (the minimum current necessary to elicit an action potential) is higher in D2+ compared to D1+ neurons (**Figure 3D**). Given the lack of differences in the AP threshold (**Figure 3G**), a higher rheobase value in D2+ neurons may arise from the summation of small (below significance threshold) differences in membrane parameters influencing the rheobase, such as membrane potential, time constant, resistance and/or capacitance.

During patch clamp recordings, cells were filled with biocytin, allowing *post-hoc* morphological reconstruction of their dendritic trees (**Figure 4A and B**). Previous data suggested possible differences in the width of the apical dendrite shafts (measured 5 µm above the soma) between D1+ and D2+ in the prefrontal cortex (24), therefore we performed similar measurements and analysis, but have not found similar differences between D1+ and D2+ M1 pyramidal neurons (**Figure 4C and D**). Due to the relatively small slice thickness (∼200 µm, which was required to detect tdTomato fluorescence with our recording setup) and resulting partial truncation of the apical dendrite tufts, only the basal dendritic trees were compared. The Sholl intersection profile revealed higher complexity of basal dendritic tree of D2+ cells (**Figure 4E**). Performed ANOVA and subsequent post hoc tests revealed that D2+ neurons have significantly more complex basal dendritic trees in the distal parts of the basal dendrites. We also found, that D2+ neurons, as compared to D1+ cells, have longer total dendritic length, a larger number of primary dendrites, and a larger number of dendritic tips (**Figure 4F**). Taken together, we found that layer V D2+ pyramidal neurons are characterized by increased intrinsic excitability and higher dendritic tree complexity.

**Figure 4.**
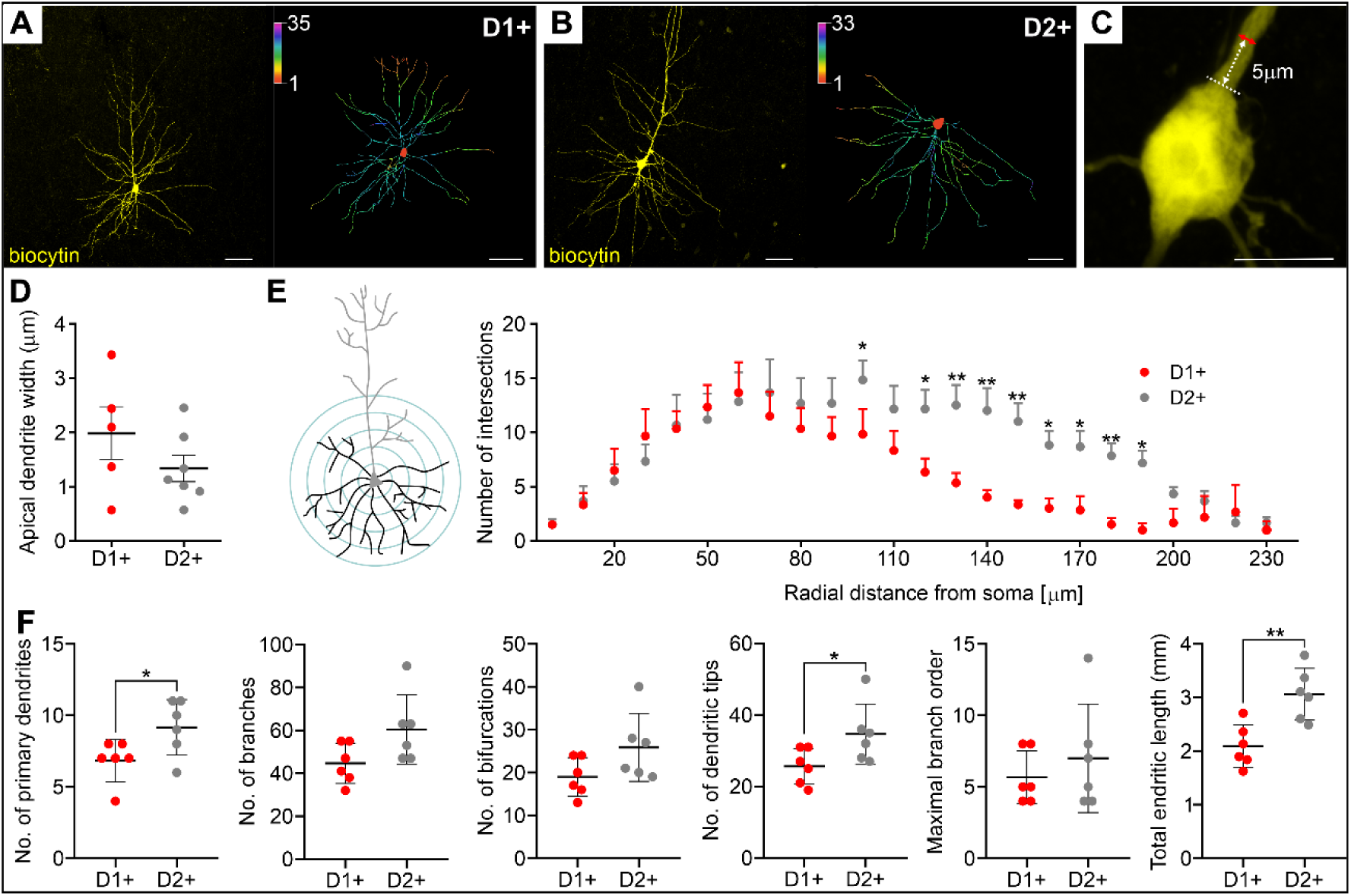
Layer V D2+ pyramidal neurons have more complex basal dendrite morphology. (**A** and **B**) Representative images of biocytin-filled layer V pyramidal D1+ and D2+ neurons (left panels) and 2D reconstruction of their basal dendrites (right panels). Scale bars: 50 µm. (**C** and **D**) Representative image of layer V pyramidal cell soma and measured widths of the shafts of D1+ and D2+ neuron dendrites. Scale bar: 10 µm. (**E**) Representation of the Sholl analysis – quantification made by superimposing a series of concentric circles of gradually increasing radius around the soma. The Sholl intersection profile, obtained by counting the number of dendritic branches at a given distance from the soma. (**F**) Morphological parameters of D1+ and D2+ neurons. (**D**-**F**) Results are displayed as mean ± SEM. ** *P* < 0.01, * *P* < 0.05, 2-way ANOVA followed by Bonferroni’s post hoc test or Student’s t-test.

### D1+ and D2+ motor cortex neurons have distinct projection targets across the brain

Motor cortex pyramidal neurons comprise three broad classes of output projection neurons: (1) intratelencephalic (IT), targeting cortical and striatal regions with somas distributed across layers II-VI; (2) pyramidal tract (PT), innervating the brainstem and spinal cord with somas located in layer V; (3) corticothalamic (CT), projecting to the thalamus with somas in layer VI (28). We used anterograde tracing to trace axonal projections of D1+ and D2+ neurons in Drd1aCre and Drd2Cre mice and observed that both D1+ and D2+ neurons originating in M1 have extensive projections to the cortical and subcortical regions of the brain. Their fibers are present in contralateral M1 (**Figure 5A**), ipsilateral somatosensory cortex (**Figure 5B**), cerebral peduncle (**Figure 5C**), pontine nuclei (**Figure 5D**), medullary pyramids (**Figure 5E**) and their decussation (**Figure 5F**). In addition, we found that D1+ neurons innervate the dorsolateral striatum (**Figure 5A**) and thalamic nuclei (**Figure 5B**). These results show, that D1+ cells constitute the majority of output projection neurons from the motor cortex and that this is a heterogenous group that overlaps with the three main classes of output neurons. In turn, D2+ neurons are a relatively smaller fraction of output projection neurons that primarily constitute fibers of corticocortical and to a lesser extent corticopontine and corticospinal tracts. Overall, the output patterns of D1+ and D2+ neurons indicate that both classes of projection neurons are well positioned to mediate the control of skilled motor behavior.

**Figure 5.**
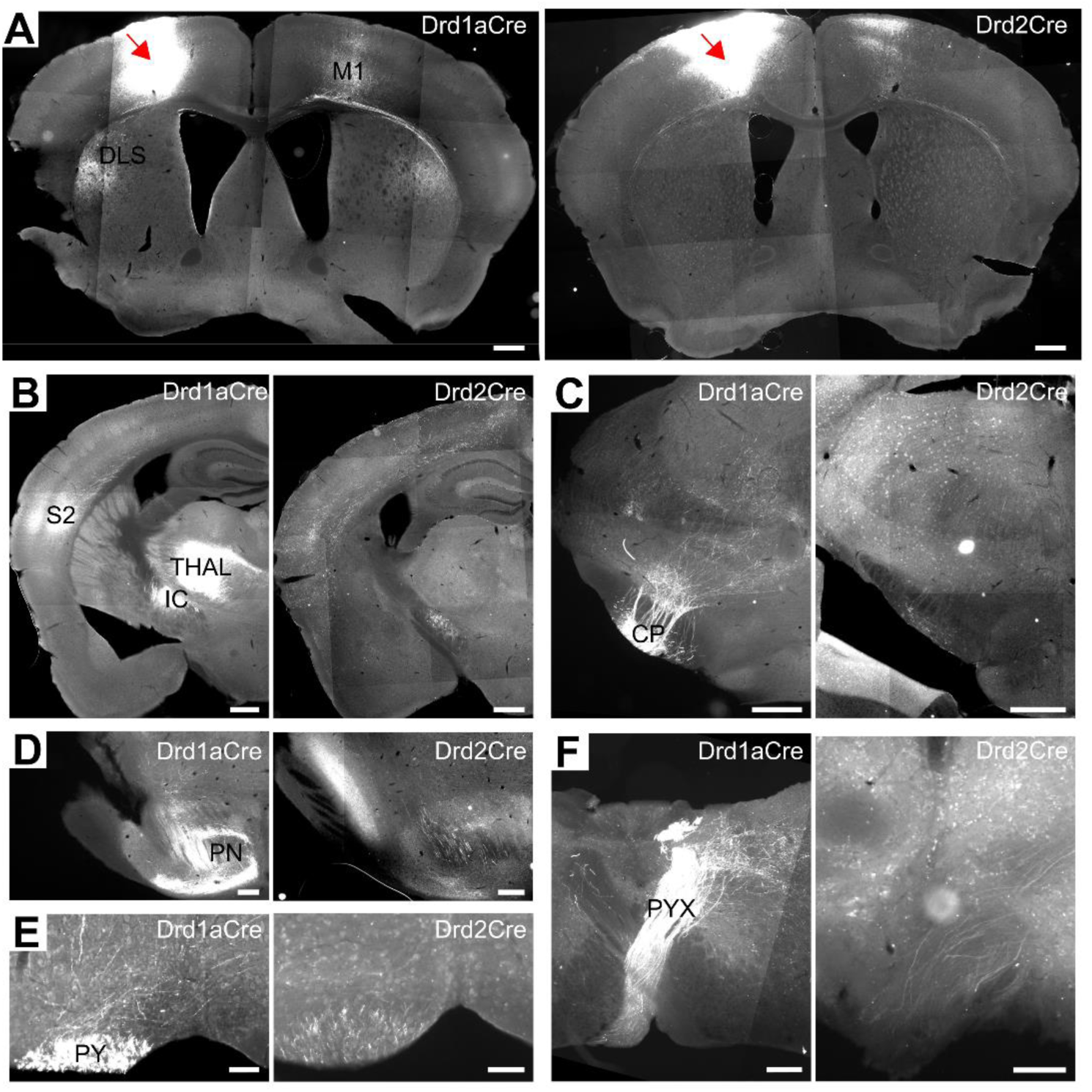
D1+ and D2+ neurons have extensive projection patterns. (**A**-**F**) Coronal sections obtained from Drd1aCre and Drd2Cre mice injected with DIO-mCherry, red arrow indicate the injection site (replicated in *n* = 3 animals from each strain). (**A**) Forebrain (0.50 from Bregma): DLS – Dorsolateral Striatum; M1 – Primary Motor Cortex. Scale bars: 500 µm. (**B**) Forebrain (-1.34 from Bregma): S2 – Secondary Somatosensory Cortex; THAL – Thalamus (Posterior and Ventral Thalamic Nuclear Groups); IC – Internal Capsule. Scale bars: 500 µm. (**C**) Midbrain (-2.70 from Bregma): CP – Cerebral Peduncle. Scale bars: 500 µm. (**D**) Pons (-4.24 from Bregma): PN – Pontine Nucleus. Scale bars: 200 µm. (**E**) Medulla (-5.80 from Bregma): PY – Pyramidal Tract. Scale bars: 100 µm. (**F**) Medulla (-8.12 from Bregma): PYX – Pyramidal Decussation. Scale bars: 200 µm (left panel), 100 µm (right panel).

### Inhibition of D2+ neurons disrupts skilled forelimb grasping

The motor cortex is involved in the control of skilled forelimb movements, particularly reaching and grasping (15). To test the functional role of newly identified classes of D1+ and D2+ neurons in controlling the execution of skilled forelimb movements, we used a chemogenetic approach (29) and the pellet-reaching task (30) (**Figure 6A and B**). Drd1aCre, Drd2Cre mice and their wild-type (WT) littermates were injected with the DIO-hM4Di virus, inducing Cre-dependent expression of G_i_-coupled designer receptors exclusively activated by designer drugs (DREADDs), in the left M1, contralateral to the reaching forelimb (**Figure 6C**). This allowed for cell type-specific inhibition of neural activity by injections of DREADDs ligand, clozapine N-oxide (CNO) (**Supplemental Figure 2**). Mice were trained to extend their right forelimb through a narrow slit to grasp and retrieve a single food pellet located on an elevated platform. Once their performance was stable across 3 consecutive days of the pre-test, mice were intraperitoneally injected with CNO (2 mg/kg). While the performance of Drd1aCre mice remained intact, we observed a reduction of successful reaches after the activation of inhibitory DREADDs in Drd2Cre mice (**Figure 6D**). A detailed analysis of individual reach outcomes revealed that the CNO-induced reduction of success rate in Drd2Cre mice was associated with degraded performance during the grasp phase, as evidenced by increased ‘no-grab’ incidence (**Figure 6E**). We further tested whether chemogenetic inhibition of D1+ or D2+ M1 neurons influences movement kinematics, but no significant effect on movement distance or speed was observed in any of the groups tested (**Figure 6F**). In addition, no effects of CNO treatment on general locomotor activity were observed, when hM4Di injected mice were tested in the open field (**Supplemental Figure 3**). Overall, these results show that D2+ neurons in M1 play a specialized role in manual dexterity, rather than gross forelimb kinematics or locomotor behavior.

**Figure 6.**
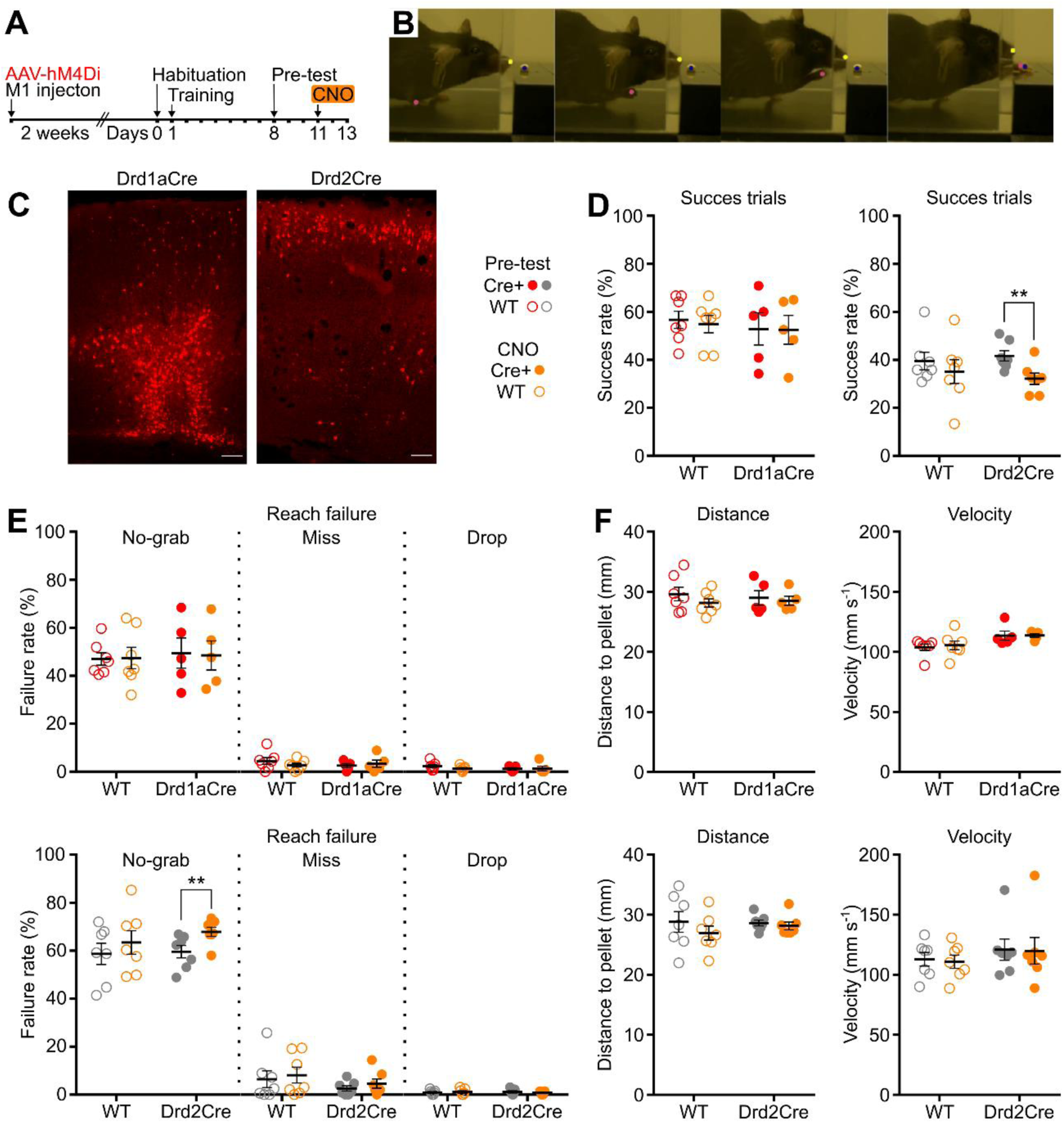
Chemogenetic inhibition of D2+ cells disrupts skilled grasping. (**A**) Timeline for the injection of inhibitory DREADDs (DIO-hM4Di-mCherry) and motor skill training experiment. (**B**) Representative images showing the pellet reaching sequence. Color circles indicate labeled body parts: nose, trained forelimb (contralateral to the inhibited motor cortex) and food pellet, which were further used for trajectory analysis. (**C**) Coronal sections showing exemplary expression of hM4Di-mCherry in D1+ and D2+ neurons in M1. Scale bars: 100 µm (**D**) Success rate expressed as a percentage of trials in which the pellet was successfully retrieved. (**E**) Reach failure represented as a percentage of reach outcomes (‘no-grab’, ‘miss’ or ‘drop’) calculated from the total number of reaching attempts. (**F)** Trajectory analysis of the first reach attempts in the trial, from the moment the paw was lifted from the ground until it touched the pellet. Kinematic variables analyzed: paw distance to pellet and velocity. (**D**-**F**) Drd1aCre *n* = 5, wild-type *n* = 7; Drd2Cre *n* = 7, wild-type *n* = 7 (WT = wild-type). Data from the respective ‘pre-test’ and ‘CNO’ (orange circles) days are collapsed across sessions and averaged to account for day-to-day variability in behavior. Results are displayed as mean ± SEM. ** *P* < 0.01, 2-way ANOVA followed by Bonferroni’s post hoc test.

## Discussion

Here, we have identified two differentially distributed, largely nonoverlapping populations of dopamine receptor-expressing neurons in the motor cortex. Specifically, we observed that neurons located in the superficial layer II/III predominantly expressed D2 receptors, while the majority of the cells located in the deep output layers V and VI expressed D1 receptors. Our observation corresponds with previous studies demonstrating the segregated expression of D1 and D2 receptors in striatal and prefrontal neurons (10–12), suggesting that this segregation is well conserved throughout the central nervous system and can reflect the existence of distinct populations of neurons with different functional roles. The majority of dopamine receptor-expressing cells in M1 were found to be glutamatergic pyramidal neurons with extensive corticocortical and subcortical projections. Interestingly, within the same output layer V, we observed significant differences in the morphological and electrophysiological properties between D2+ and D1+ pyramidal cells, with the former exhibiting increased intrinsic excitability and greater complexity of the dendritic tree. These findings are in line with previous reports showing differences in the morphology of dopamine receptor-expressing pyramidal cells located in the prefrontal cortex (24–26). Collectively, these reports and our present results, strongly suggest that the length and complexity of the dendritic tree are unique features that distinguish between D1+ and D2+ pyramidal neurons, across prefrontal and motor areas of the cerebral cortex.

Previous studies have shown that motor skill learning is accompanied by functional and structural reorganization in the upper layers of the M1 forelimb region, contralateral to the trained forelimb (31–33). This reorganization process has been shown to be dependent on dopamine signaling (19, 34), and to coincide with the emergence of population neuronal activity during learning (35). Our study revealed that inhibition of D2+ neurons in the motor cortex disrupts skilled forelimb reaching in adult mice that have learned the task. This observation aligns with previous reports demonstrating that blocking D2 receptors impairs skill learning and reduces plasticity within M1 (34). As previously discussed, D2+ neurons exhibit increased intrinsic excitability, a hallmark of the formation of neuronal ensembles (36). Therefore, D2+ neurons may be predisposed for recruitment during skill learning and subsequent reactivation during learned movement execution. Nevertheless, the selectivity of this effect was surprising, as the D1+ neurons constitute a strong representation of layer V PT neurons that also undergo rapid plastic changes during motor skill learning (37) and can directly mediate spinal circuits for skilled reaching and grasping (38). However, a recent study showed that inhibition of PT neurons does not affect reaching and grasping in mice, while inhibition of IT neurons severely disrupts performance of the skilled reach-to-grasp task (39). In line with this, D2+ neurons largely comprise the IT-type subclass of neurons located in superficial layer II/III. Considering the top-down organization of M1 (40, 41), where layer II/III neurons provide output to neurons located in lower layers, the role of D2+ neurons in motor performance may be of greater significance. Moreover, recent studies show that layer II/III neurons exhibit more prevalent direction-selective activity than layer V neurons (42, 43) and encode previous reach outcomes independent of kinematics and reward (44), which suggests that they play a superior role in motor skill learning and subsequent movement execution.

Taken together, our findings have important implications for understanding the pathophysiology of movement disorders, in particular Parkinson’s Disease. The M1 is the most important structure for volitional motor control and dopamine depletion influences the firing rate and synchronization of motor cortex neurons (45). While our study did not directly investigate the mechanisms underlying the effect of experimental PD on cortical neural activity, we did observe that inhibition of D2+ neurons in M1 results in impaired motor performance in a forelimb reaching task. This observation allows us to speculate that this neuronal population can contribute to upper limb motor impairments observed in PD patients. Experimental data showing that D2 receptor blockade-induced bradykinesia is associated with reduced motor cortex activity (46), as well as reduced neuroplasticity in the human M1 (47), support this hypothesis. Moreover, as a growing body of evidence suggests, D2 receptor activation increases the excitability of pyramidal neurons in PFC and M1 (18, 24, 48). Therefore, we propose that the therapeutic profile of neurostimulation methods aimed at modulating motor cortex activity, which have already demonstrated efficacy in improving motor symptoms of PD (49–51), could potentially be significantly augmented by cell-type specific modulation of cortical dopaminoceptive circuits, through pharmacological activation of dopamine receptors.

## Methods

### Animals

Animals were housed 2 – 5 per cage in an animal facility room with a controlled temperature (22 ± 2°C) and humidity (40-60% RH), under a 12 h light/dark cycle. Unless otherwise specified, mice had *ad libitum* access to water and laboratory chow (RM1A, Special Diet Services). Drd1aCre mice (52) were obtained from the German Cancer Research Center, Heidelberg and Drd2Cre (53) from the University of California, Davis (MMRRC_032108-UCD). Drd1a-tdTomato line 6 (54) and Ai14 (tdTomato) Cre reporter line (55) were purchased from The Jackson Laboratory (IMSR_JAX:016204 and IMSR_JAX:007914). All mice were congenic with the C57BL/6N background (>8 generations of backcrosses prior to initiation of the study). For the purpose of the project, the Drd2Cre strain was crossed with Drd1a-tdTomato and Ai14 (tdTomato) strains to obtain double transgenic animals. Genotyping was performed using a standard PCR assay according to previously described protocols and genotyping protocols available in the JAX database. The age of animals at the onset of the experiments was 8-12 weeks, with the exception of the patch clamp experiments where 6-8 weeks old animals were used. Each experimental group consisted of mice from at least 2 litters. Both sexes were used.

### Stereotaxic injections

Animals were anesthetized with a mixture of ketamine (100mg/ml) and xylazine (20mg/ml) and positioned in a stereotaxic frame (ASI Instruments). Body temperature was maintained at 37°C throughout the procedure. A small craniotomy was made above the caudal forelimb area of the motor cortex using coordinates relative to Bregma: 0.3 ± 0.2 mm anterior, 1.5 ± 0.2 mm lateral, 1.5 ± 0.2 mm ventral. Borosilicate glass pipettes with 30-40 µm tip diameters were backfilled with oil and Cre-dependent viral vectors and connected to a microliter glass Hamilton syringe via Tygon tubing. 200 nl of solution was pressure injected in batches of 20 nl, with 60 s intervals between injections. The pipette was left in the tissue for 5 min to ensure effective diffusion, and was then slowly withdrawn. After the surgery, subcutaneous injections of anti-inflammatory drug Tolfedine 4% and 5% glucose solution were given to alleviate pain and prevent dehydration. Similar procedure was followed in case of striatal injections, where Cre-dependent hM4Di viral vector was injected into dorsal striatum using coordinates relative to Bregma: 0.1 ± 0.1 mm anterior, 2.0 ± 0.1 mm lateral, 3.0 ± 0.2 mm ventral (site 1); 0.4 ± 0.1 mm anterior, 2.0 ± 0.1 mm lateral, 3.0 ± 0.2 mm ventral (site 2); 0.7 ± 0.1 mm anterior, 2.0 ± 0.1 mm lateral, 3.0 ± 0.2 mm ventral (site 3).

For the purpose of tracing experiments Drd1aCre and Drd2Cre mice were injected with rAAV2-Ef1a-DIO-mCherry and Drd2Cre::Drd1a-tdTomato mice were injected with rAAV2-Ef1a-DIO-EYFP. For behavioral experiments and multi-electrode array (MEA) recordings Drd1aCre, Drd2Cre mice and their wild-type littermates received injections of pAAV-hSyn-DIO-hM4D(G_i_)-mCherry. Viral vectors were obtained from Addgene and UNC Vector Core. Animals were returned to their home cages for 2 weeks before being used in behavioral and MEA experiments, or 4 weeks in case of anatomy and tracing studies.

### Histology and fluorescence microscopy

Mice were euthanized via intraperitoneal injection of Morbital (Biowet) (sodium pentobarbital 133.3 mg/ml + pentobarbital 26.7 mg/ml) and perfused with ice-cold phosphate-buffered saline (PBS, pH 7.4) followed by 4% formaldehyde (pH 7.4). Dissected brains were fixed in 4% formaldehyde for 12h at 4°C. Coronal slices (40-50 µm) containing the motor cortex and striatum, thalamus, midbrain, pons and medulla were cut on a VT1000S vibrating blade microtome (Leica Biosystems).

In the case of Drd1a-tdTomato and Drd2Cre::Drd1a-tdTomato mice, the tdTomato signal was enhanced with antibody against red fluorescent protein (RFP, Rockland). For tdTomato labeling, slices were first blocked in PBS-T containing 10% NDS (Jackson ImmunoResearch) and 0.6% Triton X-100 (Sigma-Aldrich) for 1 h at room temperature; followed by incubation with primary antibody anti-RFP 1:1000 (Rockland) in PBS-T (2% NDS, 0.3% Triton X-100) overnight at 4°C; and secondary antibody Cy3 1:400 (Jackson ImmunoResearch) overnight at 4°C. Finally, rinsed slices were mounted on glass slides and coverslipped with Fluoroshield containing DAPI (Sigma-Aldrich). Slices obtained from Drd1aCre, Drd2Cre or Drd2Cre::Ai14 mice were mounted on glass slides and coverslipped, without additional incubation.

Images were acquired using Axio Imager.M2 fluorescence microscope (Zeiss) equipped with Axiocam 503 mono camera. Whole brain images were acquired with 5x objective and z-stack images of regions of interest were acquired with 20x objective. ZEN software (Zeiss) was used for initial image acquisition and preprocessing. ImageJ (56) software was used for subsequent image processing that involved cropping and adjustment of brightness and contrast. Cell counting was performed in a ∼700 µm wide region of interest across the total depth of M1, using the ImageJ Cell Counter plugin. Landmarks indicating the borders between layers were identified based on the distance from the pia to the white matter, as described previously (57, 58).

### Multiplex fluorescent in situ hybridization (RNAscope)

*In situ* hybridization using the RNAscope™ HiPlex12 v2 Assay for AF488, Atto550 and Atto647 detection (Advanced Cell Diagnostics, ACD) was performed on fresh frozen 16 µm brain sections obtained from Drd1a-tdTomato and Drd2Cre::Ai14 mice as described previously (59). During the primary hybridization step the following probes were used for round 1 (R1): Drd1R (Mm- Drd1-T1, cat. no. 461901-T1, ACD), tdTomato (tdTomato-T2, cat. no. 317041-T2, ACD), Drd2 (Mm-Drd2-T3, cat. no. 406501-T3, ACD) and round 2 (R2): vGlut2 (Mm- Slc17a7-T4, cat. no. 416631-T4, ACD), vGAT1 (Mm-Slc32a1-T6, cat. no. 319191-T6, ACD). Prior to hybridization with Fluoro T1-T3 and T4-T6 the FEPE Reagent was applied to minimize tissue autofluorescence.

Images of the same M1 area for R1 and R2 were acquired with an Axio Imager M2 fluorescent microscope (Zeiss) with an automatic stage and Axiocam 503 mono camera (Zeiss), and processed using Zen (Zeiss), CorelDraw 2020 (Corel Corporation), ImageJ (56) and HiPlex Image Registration Software v1.0 (ACD).

Cells expressing the mRNAs tested were counted, using the ImageJ Cell Counter plugin, in three regions of interest: one in layer II/III and two in layers V-VI. The presence of tdTomato mRNA was used as a marker for neurons expressing either Drd1a or Drd2 receptors in Drd1a-tdTomato and Drd2Cre::Ai14 mice, respectively. Cells were identified by the presence of a cell-like distribution of fluorescent mRNA dots and/or a DAPI-stained nucleus. mRNA was assessed as present if at least two unambiguous dots were observed. The area-proportional Euler diagrams representing different cell groups and their relationships in specific layers of the cortex were generated using the open-source software Edeap (60).

### Whole-cell patch clamp recordings

Animals were deeply anesthetized with isoflurane (Baxter) and decapitated. Brains were collected in the ice-cold artificial cerebrospinal fluid (ASCF) containing (in mM); 92 NaCl, 2.5 KCl, 1.25 NaH_2_PO_4_, 30 NaHCO_3_, 20 HEPES, 5 Na+ ascorbate, 3 Na+ pyruvate, 2 thiourea, 10 glucose, 10 MgSO_4_, 0.5 CaCl_2_ (pH 7.4; osmolarity: 290-300 mOsm/kg), saturated with carbogen (95% O_2_ and 5% CO_2_).

Brains were cut into 200 µm thick coronal slices containing M1 (AP: 0.00 – 0.60 mm from Bregma) on VT1000S vibrating blade microtome (Leica). Slices were immediately transferred to the incubation chamber containing carbogenated, warm (32°C) ACSF containing (in mM): 118 NaCl, 25 NaHCO3, 3 KCl, 1.2 NaH2PO4, 2 CaCl2, 1.3 MgSO_4_ and 10 glucose. After a recovery period (90–120 min, room temperature), slices were placed in a recording chamber, where the tissue was perfused (2 ml/min) with carbogenated, warm (32°C) ACSF of the same composition. Borosilicate glass pipettes (Sutter Instruments), 6-8 Ω tip resistance, were filled with a solution containing (in mM): 145 potassium gluconate, 2 MgCl_2_, 4 Na_2_ATP, 0.4 Na_3_GTP, 1 EGTA, 10 HEPES (pH 7.3; osmolarity: 290-300 mOsm/kg) and 0.05% biocytin for subsequent immunofluorescent identification. All reagents were purchased from Sigma-Aldrich, except biocytin (Tocris Bioscience). The calculated liquid junction potential was +15mV and analyzed data were corrected for this value.

Whole-cell current- and voltage-clamp recordings were obtained from pyramidal cells located in the motor cortex deep output layer V. Cells were identified with the Axio Examiner.A1 microscope (Zeiss) using video-enhanced infrared-differential interference contrast. The neuronal identity of dopaminoceptive neurons was confirmed by the presence of tdTomato excited at 530 nm with a Colibri LED light fluorescent system (Zeiss). Signal recordings were made with a SEC-05X amplifier (NPI Electronics) and Micro3 1401 converter (Cambridge Electronic Design). The recorded signal was lowpass filtered at 3 kHz and digitized at 20 kHz.

Electrophysiological properties of examined neurons were assessed based on their responses to a series of stimulation protocols in voltage and current clamp mode. To measure the I–V relationships of the steady state current, voltage steps stimulations ranging from − 120 to - 50 mV (10 mV change, pulse duration 500 ms) were delivered from a holding potential of -75 mV. Neuronal excitability (input–output relationship) was measured as the number of spikes elicited by applied current pulses from -250 to +500 pA (50 pA increment, pulse duration 500 ms). Linear regression for excitability and steady-state current I–V relationship was calculated in GraphPad Prism software. The voltage sag was measured from the voltage response to a -250 pA hyperpolarizing current step (500 ms). The rheobase (the current value at the moment of reaching the AP threshold) was determined based on the voltage response to current ramp stimulation (0– 1 nA, 1 s). Passive membrane properties (resistance, capacitance, time constant) were calculated from the voltage responses to a -50 pA current pulse (500 ms). Action potential (AP) properties were calculated from single AP evoked from a membrane potential of - 75 mV, by a 0.5 ms depolarizing current pulse. Custom-written scripts in Signal and Spike2 software (CED) were used for data analysis.

### Morphological reconstruction

Slices from patch clamp experiments were fixed overnight at 4°C in 4% formaldehyde in PBS; blocked for 24 h at 4°C in PBS containing 10% NDS (Jackson ImmunoResearch) and 0.6% Triton X-100 (Sigma-Aldrich); followed by incubation with ExtrAvidin-Cy3 1:200 (Sigma-Aldrich) in PBS (2% NDS, 0.3% Triton X-100) for another 48 h at 4°C. Rinsed slices were mounted on glass slides and coverslipped with Fluoroshield containing DAPI (Sigma-Aldrich). To compare the dendritic morphology of recorded neurons, cells well-filled with biocytin that had clearly visible dendritic trees and no major truncations, were further imaged using a LSM 710 META laser scanning confocal microscope (Zeiss) under 20x magnification and z-stack images were taken, using LSM 780 META, under 40x magnification. Dendrite tracing and 3D reconstruction (10 µm steps) were performed in ImageJ using the Simple Neurite Tracer plugin (61). The width of the apical dendritic shaft was measured 5 µm above the soma, but due to the truncation of apical shafts in the majority of imaged cells, only the basal part of the dendritic tree was traced. L-Measure software (62) was further used, to acquire morphological parameters, e.g. total dendritic length, maximal branch order, number of primary dendrites, branches, bifurcations, and dendritic tips.

### Multi-electrode array (MEA) recordings

The multielectrode extracellular *ex vivo* experiments were performed as previously described (63). 2 weeks after the viral vector transfection (described above) mice were decapitated, and the slices were prepared as for the patch-clamp recordings. The brains were collected in carbogenated, ice-cold ACSF, comprising (in mM): 25 NaHCO3, 3 KCl, 1.4 Na2HPO4, 2 CaCl2, 10 MgCl2·6H2O, 10 glucose, 185 sucrose, and the 250 µm thick coronal slices were cut using a microtome (Leica). Sections containing striatum were transferred to an incubation chamber filled with carbogenated, preheated (32 °C) ACSF, with the same composition as for the patch-clamp recordings, with the addition of 0.01 g L−1 phenol red. The slices were incubated for 90 min and transferred to the recording wells of the MEA2100-System (Multi Channel Systems GmbH, Reutlingen, Germany). The striatum was positioned upon the recording electrodes of the 8 × 8 perforated multi-electrode array (MEA, 200 µm spacing, 60PedotpMEA200-30iR-Au, Multi Channel Systems). Carbogenated, warm (32 °C) ACSF was used for continuous perfusion of the slices (2 ml/min). Before the recording, slices were left to settle for an hour. The Clozapine N-oxide (CNO; Sigma-Aldrich) was freshly diluted in the recording ACSF (10 µM; 10 ml) and delivered by bath perfusion. The raw signal was sampled at 20 kHz and recorded using Multi Channel Experimenter (Multi Channel Systems).

The data files were converted to HDF5 and CED-64 formats using Multi Channel Data Manager (Multi Channel Systems). Using custom-written Matlab scripts and the KiloSort algorithm (64) the initial automatic spike sorting was performed. Then, the signal was transferred into the band-pass filtered (Butterworth filter; fourth order; cut off: 0.3 – 7.5 kHz) CED-64 file and single units were manually refined using principal components analysis and autocorrelation. Afterwards, the data were binned (30 s) and analyzed using NeuroExplorer 5 and the temporal heatmaps encoding the single-unit activity were prepared using a custom-written Matlab script to illustrate the drug-induced changes. A response to CNO application was considered significant if it differed by one standard deviation from the baseline mean of the recorded signal.

### Motor skill training

To evaluate the function of cortical dopamine receptor-expressing neurons, a single pellet-reaching task was performed. The design of the Plexiglas training chamber and the experimental protocol was based on the previously described protocol (30), with modifications. To ensure motivation during training, animals were food deprived to ∼85% of their initial body weight. To habituate mice to food reward, food pellets (20 mg, dustless precision pellets, Bioserv) were given in the home cage 3 days before training. To habituate mice to the testing environment, a day before training each mouse was individually placed for 20 min in the Plexiglas testing chamber (20 cm long, 8 cm wide, 20 cm high, with a 0.5 cm wide vertical slit on the front narrow wall), and was allowed to explore the apparatus, consume pellets laying on the floor or reach for multiple pellets located outside the box. During the subsequent training (days 1-7) mice were trained to extend their right forelimb (contralateral to hM4Di injected hemisphere) through a narrow slit to grasp and retrieve a single food pellet located on an elevated platform (1,5 cm high, 1 cm away from the opening and centered 0,5 cm to the left). Following this initial training, mice performance was assessed for additional 3 sessions (days 8-10) during which baseline success rate was measured (pre-test). On the subsequent test days (days 11-13), animals received an i.p. injections of clozapine N-oxide (CNO, 2 mg/kg) 30 min before being placed in the testing apparatus.

Reaching events during pre-test and CNO treatment sessions were recorded at 100 frames-per-second and 640 x 480 pixels by a color video camera (Basler acA1440-220uc). Each session consisted of 40 trials that started when the animal approached the pellet and the reach attempt was recorded for 5 s. The success rate was calculated as a percentage of trials in which the pellet was successfully retrieved (regardless of the number of reaching attempts). Animals that failed to maintain at least 30% of success rate during the pre-test were excluded from the analysis. Individual reach outcomes (from all reaching attempts) were analyzed and reach failures were classified as ‘no-grab’ (pellet was touched but not correctly grasped), ‘miss’ (pellet was not touched) or ‘drop’ (pellet was retrieved and dropped before being placed in the mouth). Attempts to reach with the tongue or left forelimb (ipsilateral to hM4Di injected hemisphere) were omitted from the analysis. To account for day-to-day variability in behavior, data from the respective pre-test and CNO treatment test days were averaged.

### Kinematic analysis

To quantify reaching kinematics we analyzed the first reach attempt (irrespective of the outcome) extracted from trials recorded in the pre-test and test sessions. The position of the paw and the pellet were tracked with DeepLabCut (65) from the moment the paw was lifted from the ground until it touched the pellet. Trials in which the starting position of the paw was not on the ground or in which the entire reaching motion was not clearly visible were excluded from analysis. In a small fraction of frames, the deep neural network failed to correctly recognize the position of the paw. In such cases, paw coordinates were linearly extrapolated. Paw distance to pellet was calculated as sqrt(x^2^ + y^2^) normalized to the number of pixels by 1 mm and velocity was calculated as a derivative of this distance.

### Open field

To assess the effects of clozapine N-oxide on basic locomotor activity, animals were tested in the open field (35,5 cm long x 25,5 cm wide x 22 cm high). Animals received an i.p. injection of CNO (2 mg/kg) 30 min prior to the placement in the arena filled with the home cage bedding. A single 10 min session was recorded at 60 frames-per-second and 640 x 480 pixels by a monochrome video camera (Creative HD 720p). The position of the animal was tracked with DeepLabCut. The total distance traveled and average movement speed were measured.

### Statistics

Results are presented as mean ± SEM. Statistical analysis was based on the assumption that the samples follow a Gaussian distribution. Student’s t-test was applied for statistical comparisons between two groups and 2-way ANOVA followed by post hoc analysis (Bonferroni’s multiple comparison test) was used for analysis with multiple groups and repeated measures were incorporated when appropriate. *P* < 0.05 was considered statistically significant. All statistical analyses were conducted using GraphPad Prism software.

### Study approval

Procedures were approved by the 2nd Local Institutional Animal Care and Use Committee in Krakow (approval number 65/2019, issued on March 07, 2019) and conducted in accordance with the directive 2010/63/EU of the European Parliament and of the Council of September 22, 2010 on the protection of animals used for scientific purposes, and with the Polish Act on the Protection of Animals Used for Scientific or Educational Purposes of January 15, 2015.

### Data availability

Any additional information required to reanalyze the data reported in this paper is available from the lead contact upon request.

## Author contributions

PEC conceived the project. PEC and AB were responsible for the overall study design. PEC performed stereotaxic injections of viral vectors and conducted behavioral experiments. PEC and MG performed histological and kinematic analyses. SD performed whole-cell patch clamp recordings with supervision from AB, morphological reconstruction of recorded cells with supervision from AG, and subsequent data analysis.

AG and AT performed and analyzed RNA *in situ* experiments. AT and KP performed MEA recordings and data analysis. LS was responsible for genotyping and maintaining the colony of transgenic animals. GK provided the Drd1aCre line and JRP provided all other transgenic mouse lines used in the study. The main part of the manuscript was written by PEC. All authors discussed the results and commented on the manuscript.

## Acknowledgments

This study was supported by the National Science Center grant SONATINA 2018/28/C/NZ4/00102 (to PEC). We thank Monika Baginska for excellent technical assistance in genotyping and maintaining the colony of transgenic animals.

