## Supplemental Figure 1 for "Dopamine receptor-expressing neurons are differently distributed throughout layers of the motor cortex to control dexterity"

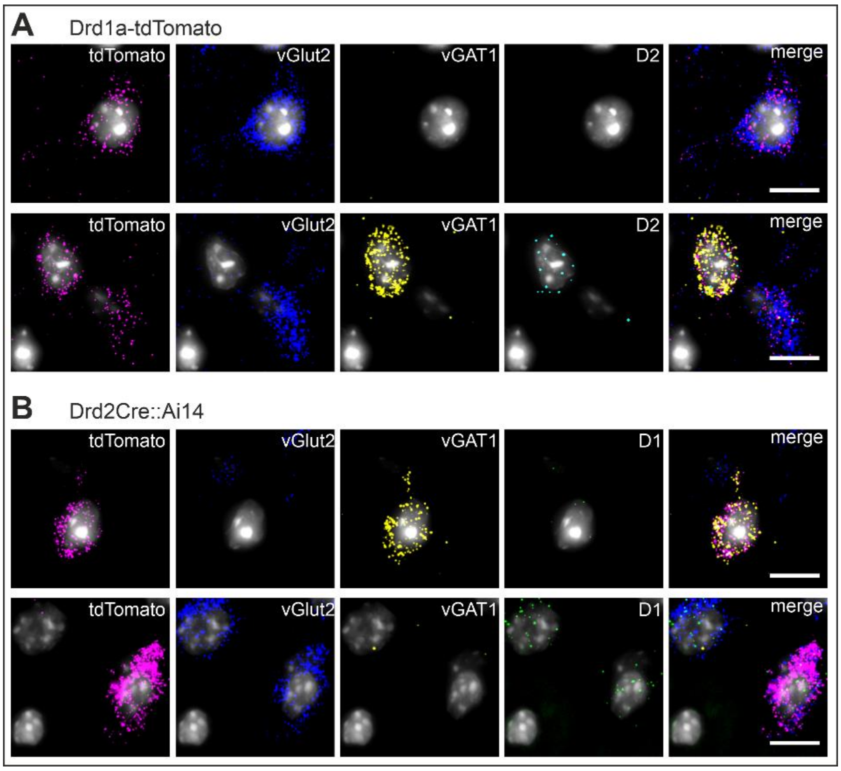


**Supplemental Figure 1| Drd2 and Drd1a mRNA co-localization in tdTomato+ cells in Drd1a-tdTomato and Drd2Cre::Ai14 mice.**

(**A and B**) Representative images showing co-localization of tdTomato (pink) mRNA and either vGlut2 (blue) or vGAT1 (yellow) mRNAs, and either Drd1a (green) or Drd2 (turquoise) mRNAs within an individual neuron. Scale bar: 10 µm.
