## Supplemental Figure 2 for "Dopamine receptor-expressing neurons are differently distributed throughout layers of the motor cortex to control dexterity"

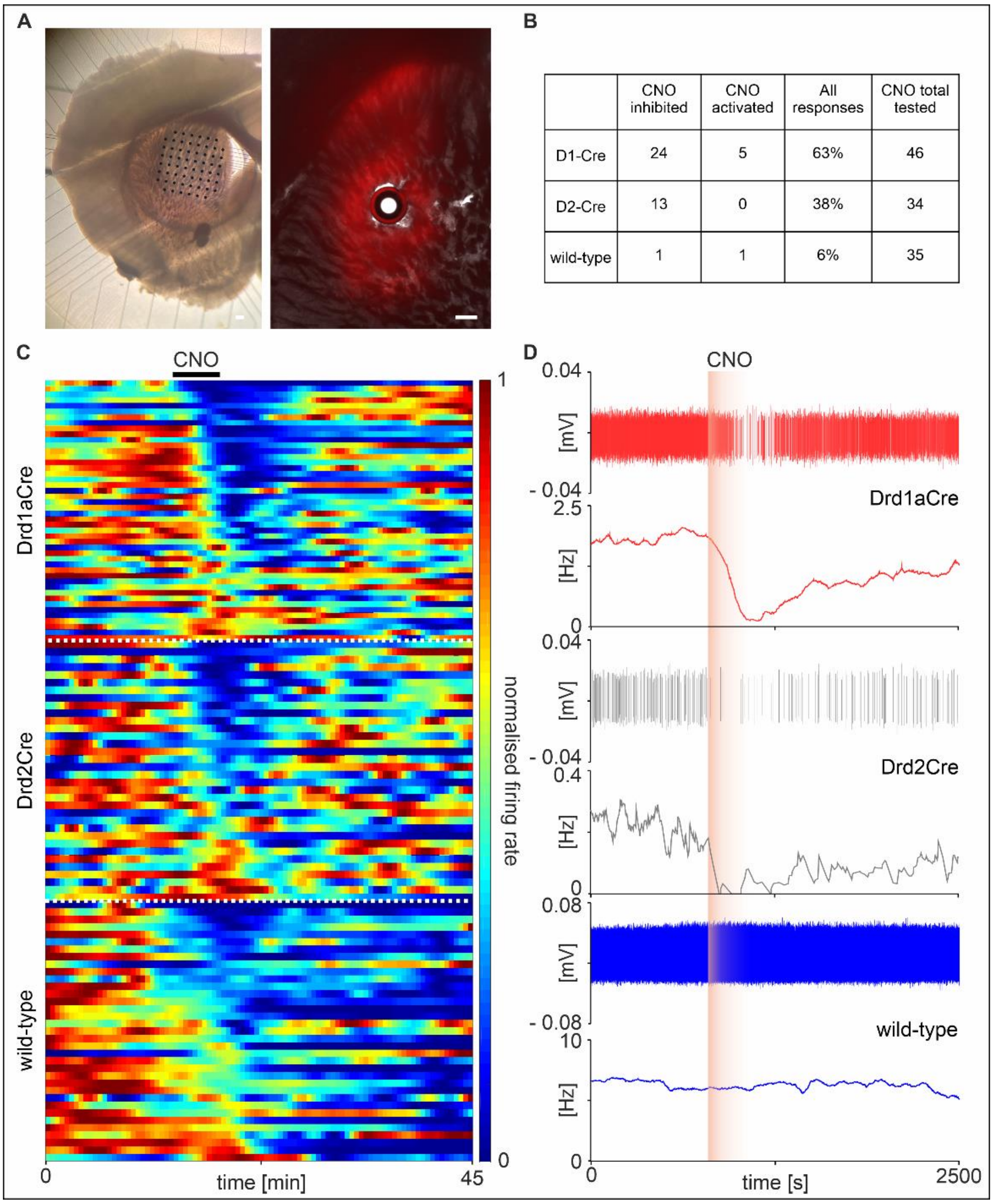


| | CNO inhibited | CNO activated | All responses | CNO total tested |
|---|---|---|---|---|
| D1-Cre | 24 | 5 | 63% | 46 |
| D2-Cre | 13 | 0 | 38% | 34 |
| wild-type | 1 | 1 | 6% | 35 |

**Supplemental Figure 2| CNO-induced inhibition of hM4Di expressing striatal projection neurons in Drd1aCre and Drd2Cre mice.**

(**A**) Coronal section obtained from a wild-type (left) and Drd1aCre mouse (right) showing slice placement upon the multi-electrode array (black dots indicate single recording electrodes) and hM4Di-mCherry expression in the striatum after recording. Scale bars, 200 µm. (**B**) Table summarizing recorded neurons and their responses to CNO application. (**C**) Temporal heatmaps encoding single-unit activity (SUA) of all recorded neurons and their response to CNO application (10 µM, 10 ml; indicated by a black line).

Each row indicates a single unit. White dotted horizontal lines classify units to either Drd1aCre mice (*n* = 46 units), Drd2Cre mice (*n* = 34), or wild-type mice (*n* = 35). The cell activity was normalized from 0 to 1 and was sorted by the defined window corresponding to the CNO response. Bin, 30 s. (**D**) Examples of neurons recorded during CNO administration in Drd1aCre, Drd2Cre and wild-type mice. Top panels show separated spikes of a single neuron, bottom panels show a corresponding frequency histogram. The orange-shaded rectangle indicates the duration of CNO action. Bin, 60 s. (**B**-**D**) Slices were obtained from *n* = 1 hM4Di-mCherry injected animal for each group, *n* = 4 slices per animal.
