## Supplemental Figure 3 for "Dopamine receptor-expressing neurons are differently distributed throughout layers of the motor cortex to control dexterity"

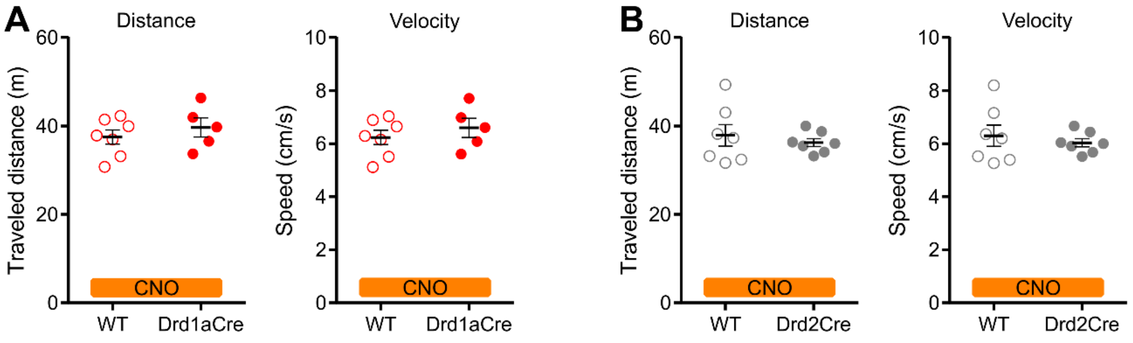


**Supplemental Figure 3| Locomotor activity in the Open Field is not altered by chemogenetic inhibition of D1+ or D2+ cells.**

**(A and B)** Analysis of the distance traveled and velocity during the exploration of the Open Field. The same animals were used as in the motor skill training. Drd1aCre $n$ = 5, wild-type $n$ = 7; Drd2Cre $n$ = 7, wild-type $n$ = 7 (WT = wild-type). Results are displayed as mean ± SEM.
