## Supplemental Table 1 for "Dopamine receptor-expressing neurons are differently distributed throughout layers of the motor cortex to control dexterity"

**Supplemental Table 1.** Combinations of mRNA species detected in M1 slices obtained from Drd1a-tdTomato and Drd2Cre::Ai14 mice.

| mRNA Combination | Drd1a-tdTomato | | Drd2Cre::Ai14 | |
|---|---|---|---|---|
| | Layer II-III [Mean ± SD (%)] | Layers V-VI [Mean ± SD (%)] | Layer II-III [Mean ± SD (%)] | Layers V-VI [Mean ± SD (%)] |
| All vGlut2 | 62 ± 20 (100) | 52 ± 14 (100) | 67 ± 12 (100) | 48 ± 11 (100) |
| vGlut2 only | 53 ± 18 (85) | 12 ± 5 (23) | 42 ± 10 (63) | 44 ± 10 (92) |
| vGlut2/tdTomato | 9 ± 10 (15) | 40 ± 16 (77) | 25 ± 6 (37) | 4 ± 2 (8) |
| All vGAT1 | 10 ± 1 (100) | 8 ± 5 (100) | 12 ± 4 (100) | 9 ± 4 (100 |
| vGAT1 only | 3 ± 2 (30) | 2 ± 2 (25) | 8 ± 3 (67) | 5 ± 2 (56) |
| vGAT1/tdTomato | 7 ± 2 (70) | 6 ± 3 (75) | 4 ± 1 (33) | 4 ± 3 (44) |
| All vGlut2/vGAT1 | 1 ± 1 (100) | 0 (0) | 2 ± 2 (100) | 1 ± 1 (100) |
| vGlut2/vGAT1 only | 0 (0) | 0 (0) | 1 ± 1 (50) | 1 ± 1 (100) |
| vGlut2/vGAT1/tdTomato | 1 ± 1 (100) | 0 (0) | 1 ± 1 (50) | 0 (0) |
