## Supplemental Table 2 for "Dopamine receptor-expressing neurons are differently distributed throughout layers of the motor cortex to control dexterity"

**Supplemental Table 2.** Combinations of Drd1a and Drd2 mRNA species detected in tdTomato+ neurons in Drd1a-tdTomato and Drd2Cre::Ai14 mice.

| mRNA Combination | Drd1a-tdTomato | | Drd2Cre::Ai14 | |
|---|---|---|---|---|
| | Layer II-III [Mean ± SD (%)] | Layers V-VI [Mean ± SD (%)] | Layer II-III [Mean ± SD (%)] | Layers V-VI [Mean ± SD (%)] |
| All tdTomato | 18 ± 10 (100) | 46 ± 15 (100) | 30 ± 5 (100) | 9 ± 4 (100) |
| tdTomato only | 15 ± 11 (83) | 44 ± 15 (95) | 29 ± 6 (96) | 4 ± 2 (39) |
| tdTomato/D2 | 3 ± 2 (17) | 2 ± 1 (5) | - | - |
| tdTomato/D1 | - | - | 1 ± 2 (4) | 6 ± 3 (61) |
